# A Strategy Study on Risk Communication of Pandemic Influenza——A Mental Model Study of College Students in Beijing

**DOI:** 10.1101/490532

**Authors:** Honglin Yang, Linxian Wang, Yadong Wang, Bo Zheng, Shuai Du, Xinyi Lu

**Author notes:** Email address (Honglin Yang), (LinXian Wang) (Yadong Wang), (Bo Zheng), (Shuai Du), (Xinyi Lu).

## Abstract

Understanding the risk perception of pandemic influenza can improve the risk communication efficiency of the government and ultimately reduce losses caused by the disaster. A mental model interview of 28 individuals who discussed pandemic influenza was analyzed in this paper. The interviewees were college students in Beijing, China who were evaluated to understand their views on the risk perception of pandemic influenza. Referring to the mental model theory, the researchers using Delphi method to identify the key risk factors and concepts to examine the public understanding of these contents; then, the researchers identify the deviations in their understanding so that suggestions and countermeasures have been put forward to enhance the effectiveness of risk communication. Most of the conceptual content was mentioned by most interviewees. However, some interviewees showed misunderstanding including excessive optimism about the consequences of pandemic influenza, a lack of detailed mitigation measures, and negative attitudes toward health education and vaccination. Once faced with threats, this may lead to the failure of risk communication. In Beijing City, the center of domestic and international education, the historical SARS epidemic and this year’s seasonal flu peak are all hints of the potential risk of a pandemic outbreak. Beijing’s college students’ one-sided understanding and misunderstanding of the relevant risk information may increase the risk during an influenza pandemic. The results highlight the necessity for the government to clearly focus on the communication content of the student group, provide an official reference plan for the public and update health education on this topic.

## Introduction

Influenza is a highly variable infectious disease that can easily evolve into a pandemic and pose a major threat to people’s lives and safety. (1). The corresponding emergency response measures require the active cooperation of the public to produce an effective response. Because of its wide range of impact and potential mortality, effective risk communication will help the public understand the information related to influenza (2). But in fact, our risk communication and health education are still lacking. In January 2018, an article from the micro-blog, named “the middle aged people under Beijing flu”, has attracted wide attention. The author, a college student, shared his noticeable blog article which records how his grandfather was infected with influenza and passed away. The article included the whole process of his family’s infection and coping with influenza. However, their decisions are mostly erroneous and misleading, because a lacking of understanding of influenza. These issues further illustrate the urgent need to effectively communicate the risk of pandemic influenza to general public in china.

When public health emergencies occur, the government often asks experts what the public should know; thus, how to effectively transform scientific knowledge into a concept that can be understood by a useful structure and a nonprofessional background is a key component of an effective risk communication officer (3). Mental Model Theory holds that people’s understanding of information and judgment of decision-making are influenced by their own unique mental models. The mental model is a causal belief (3,9,10) that guides the decisions and actions of the world. A person’s mental model consists of many factors, including personal experience, acquired learning, analogy and reasoning (3,11,12,13,14,15), which could be a complex system in our mind. The mental model analysis, which combines a qualitative research method of cognitive psychology and communication, shows the cognition of two different groups to the same thing by the form of mind map. It’s convenient for us to compare the cognitive differences and characteristics between the two groups in the same thing. For experts and the public, if we can find the deviations in risk perception, we can formulate corresponding communication strategies to correct these deviations. Thus, targeted health education can help one modify their mental model to establish a sense of risk and improve the risk management of influenza. What’s more, in China there is no normalized and systematic health education or publicity related to large-scale infectious diseases for general public. Until there is an outbreak of infectious diseases, there will be short-term targeted experts recommend to the public. And compared with other places, colleges and universities as a large number of students concentrated in the key areas, personal health literacy and living habits vary greatly, once the campus flu patients, the risk of large-scale infections is extremely high, and produce a multi-faceted serious consequence. For college students who need to live independently for a long time, it is necessary to study and take care of their personal life at the same time. It is necessary to carry out targeted health education and learn some basic knowledge of influenza pandemic prevention. Our study uses a mental model analysis to find out the cognitive characteristics of college students’ risk of pandemic influenza by comparing each student’s understanding of expert knowledge and diagnosing which information can improve their decision-making and thus their security (4,5,6,7,8). When a pandemic threatens, proper risk communication will reduce losses. The purpose of this paper is to study the key communication content of students in the influenza pandemic, hoping to provide some reference for the formulation of special emergency plans and health education materials.

Our data are based on personal interviews with 28 students from 5 universities in Beijing. This study used an open personal interview structure to learn about students’ beliefs regarding the risk of pandemics, to explore how people understand the flu and its risks, to identify the reasons for these beliefs and behaviors and to determine views on current risk communication. It is necessary for people to have knowledge to be able to protect themselves from the threat of flu. The content of this study included the degree of knowledge of the experts’ risk perception and their understanding of pandemic influenza. It is not necessary for members of the public to know all the details of influenza risk information to make wise decisions.

In China, the mental model theory has primarily been applied to team management. There has not yet been any application in the field of health education. Only risk management strategy has been studied from the perspective of public demand. Furthermore, there is no official pandemic preparedness plan for the general public. Consequently, several studies have been performed in this field. Lazrus and others have studied the public mountain flood communication framework in Boulder County, Colorado State (16). Wagner (17) found that the Alps of Bavaria influence the behavior of local floods, and to understand the local environment and conditions, more information on the floods was dependent on previous sources of experience. In the reality, many people still do not understand how real events occurred and managed, such as the 2004 Hurricane Katrina in the United States and the SARS epidemic in China in 2008, even though the government departments responded and disposed of the disaster actively. These findings reveal how the public understands risk information and makes decisions in the face of emergencies. However, most of the topics focus on natural disasters. There is no research on the application of this method in public health risk communication. The researchers hope to use pandemic influenza as an example to see early warning and emergency decision-making in a broader perspective and to determine the degree of understanding and cognition of the risk of pandemic influenza among Chinese college students to aid with government contingency plans and health education materials. This article focuses on those topics.

## Materials and Methods

### 1. Communication framework and key concepts

To succeed in risk communication, we must have a full understanding of the emergencies and identify relevant risk factors to determine the necessary communication content for general public, and we need to understand which concepts should be grasped by the public. Our study refers to the expert model (flash flood early warning system, FFW model) (18) formed by Morss et al in their study about disaster risk communication in Boulder and we modified the content which is irrelevant to pandemic flu. After that, we organized an expert seminar to revise the framework. Thus, a framework was initially formed, and the researchers summarized the concept of the corresponding framework and initially formed a communication content suitable for the influenza epidemic to adapt to the actual situation in China, including the cause of pandemic influenza, emergency preparedness and strategy, risk information and emergency response for different groups. The basic framework is shown in Figure 1. The arrow in the graph represents the interrelationship between the factors in the framework.

**Fig. 1.**
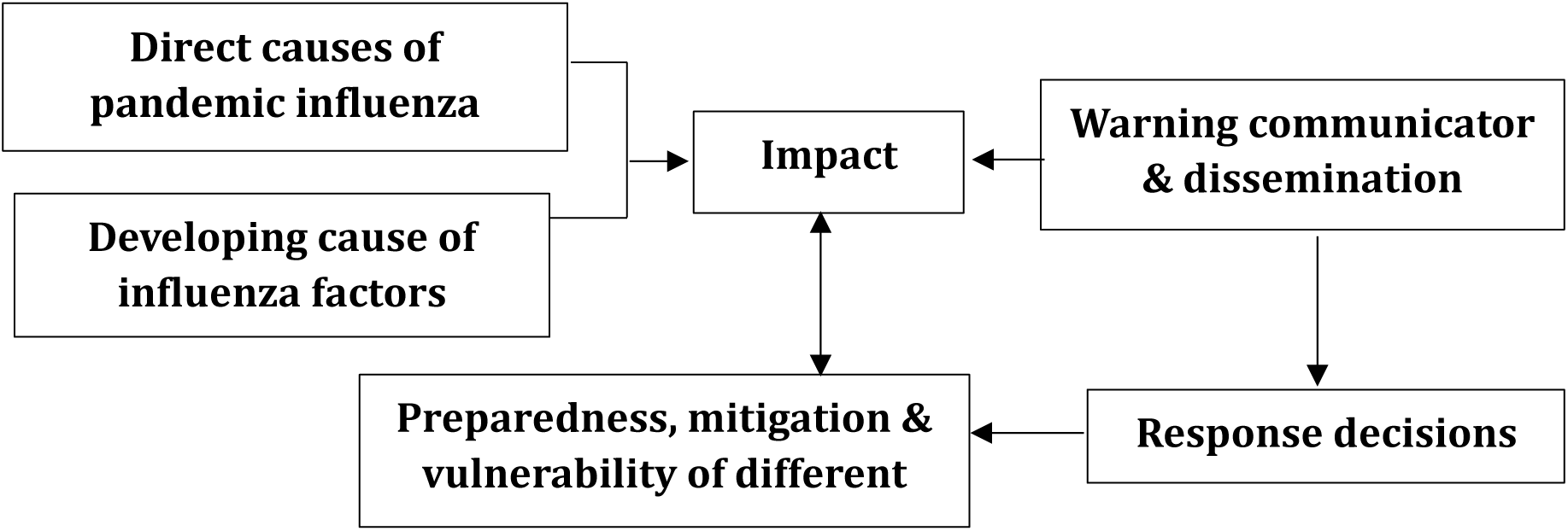
Basic framework of standardized communication for influenza: risk factors. The six boxes in the figure represent the key risk factors that make up a pandemic, and the arrows represent the relationships among the factors.

In Figure 1, the interaction between direct factors and motivation to promote development leads to the occurrence of pandemic events; the content of communication work is divided into early warning communicators, response decisions and information sources; and mitigation and emergency preparedness includes public action. In this section, different boxes represent the risk factors associated with pandemic influenza.

Next, the researchers performed a literature search and conducted expert seminars in accordance with the contents of the framework, completing the key concepts contained in each part. Finally, we used the Delphi method to amend the content of communication.

### 2. Sample and interview content

Purpose of mental model interviews is to find out which concepts or beliefs, are “out there” with some reasonable frequency then smaller samples become reasonable. There is no standard method for determining sample size in relevant theories and research practice (3). According to Professor Morgan’s monograph and related research examples, the sample size for a mental model interview should be 20∼30, at which time new information has reached saturation (3). Based on the above research facts, combined with the research design of Lazrus and Morss, the quota sampling method was selected in this study, and the first group of sample size was estimated to be approximately 30 (16,18). Through advertising poster, the researchers recruited 28 interviewees from 5 different universities. In the process of the interview, we will count the frequency of the concepts provided by the interviewees and plot a graph. We will consider increasing the sample until the respondents can no longer provide new information (3). In Figure 2, the frequency of concepts reaches its peak at No.21 interviewee. After that, the amount of information provided by respondents began to decline, which means that it was unlikely that new concept would emerge.

**Fig. 2.**
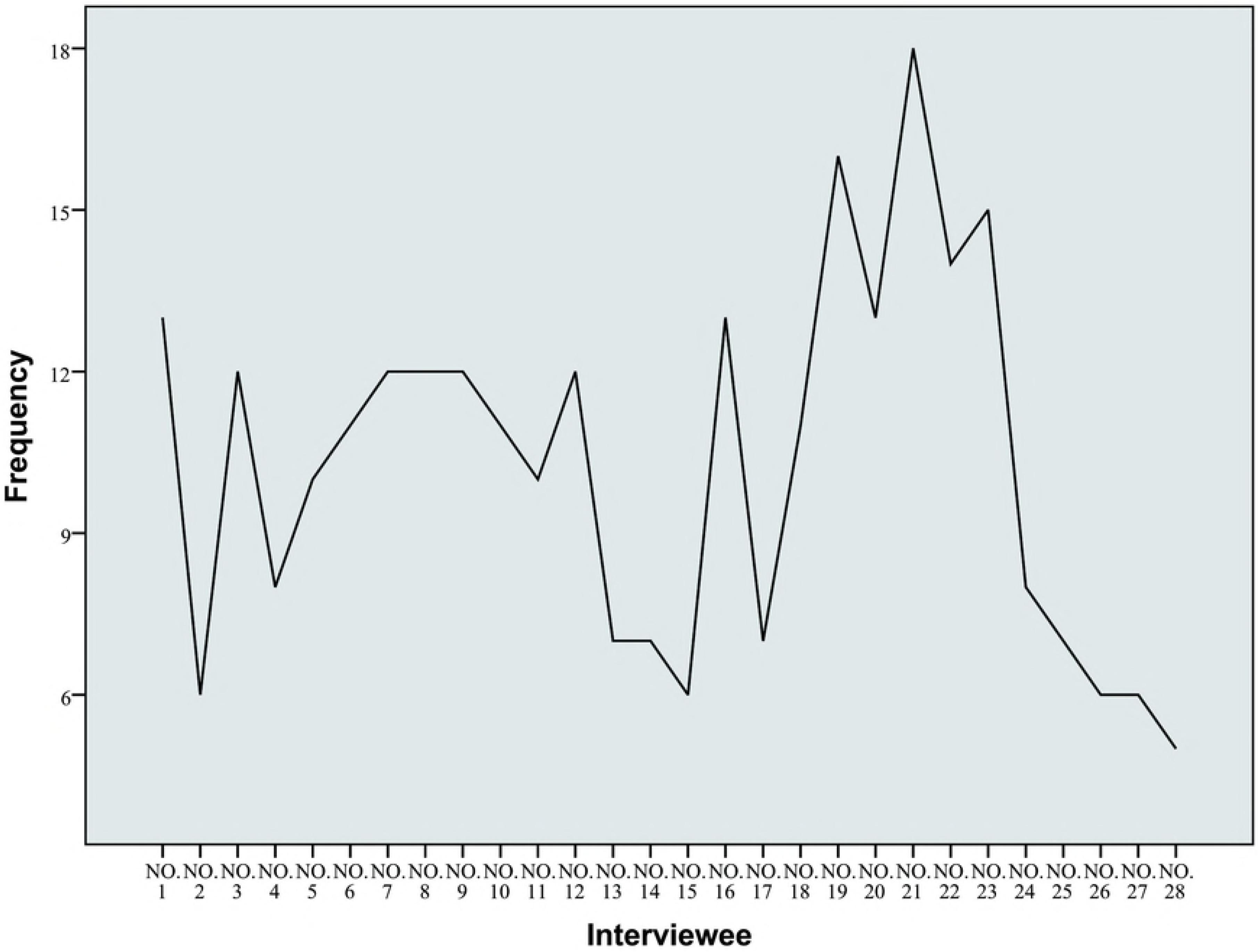
Information saturation trend provided by 28 interviewees. The fold line in the figure shows the increasing and decreasing trend of information provided by the interviewee. It is formed by connecting the points of concept frequency in Fig.3 described in the responses of 28 respondents.

The interview is face-to-face and began with an open question, such as “please tell us about the pandemic”. Because this is a public issue, the investigators guided the interviewees to elaborate on their main concepts, followed by providing details of the outbreak, including factors that may lead to a flu, the impact of that. And what mitigation measures should be taken. If the interviewer had experienced any emergencies, they were encouraged to talk about the decision or idea at that time. Finally, the interviewees were asked to make recommendations on the current status of health sector risk communication. After that, the interview results were transcribed, encoded and classified by coding software ATLAS. ti. As long as the interviewee mentioned it, a concept will be compiled, no matter how it is understood. Then, we conducted a quantitative analysis of the results of the compilation, created a statistical chart, observed the degree of attention of the interviewees, and compared those results with risk perception of experts to determine the interviewee’s understanding of the related concepts and other features.

The questions used in this interview refer to a questionnaire in the study of T. Chen and M. Haklay et al (19). The interview covers the risk factors of the pandemic, the possible impact, the emergency response including the early warning decision, and the individual’s experience of the pandemic. The results of the interviews were coded simultaneously by 2 researchers. Then, the classification consistency index (Holsti reliability) of the coder was calculated (20), which fluctuated between 0.624-0.965, and the average reliability statistic was 0.749. According to the study of Boyatzis and Burrus, the coding reliability of trained different coders ranges from 0.74-0.80 (21); therefore, the reliability of the coder was within the normal range and displayed good consistency.

## Results

### 1. Expert consultation results

**Table 1.**
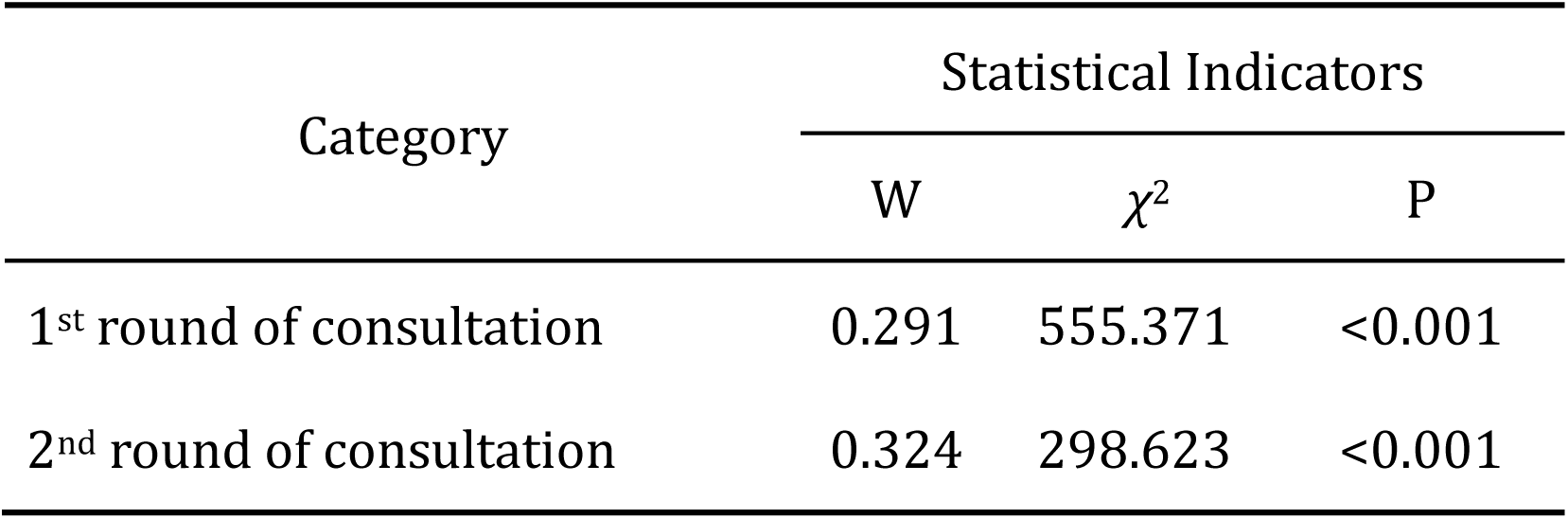
Coordination coefficient of expert consultation.

| Category | Statistical Indicators |  |  |
| --- | --- | --- | --- |
| | W | $\chi^2$ | P |
| 1<sup>st</sup> round of consultation | 0.291 | 555.371 | <0.001 |
| 2<sup>nd</sup> round of consultation | 0.324 | 298.623 | <0.001 |

There were 2 rounds of expert consultation, with a feedback rate of 100% per round of questionnaires. The 18 experts who participated in the questionnaire investigation were from the National Health Bureau, the national centers for disease control and prevention (CDC), the Beijing CDC, the health education center and the Provincial Health Bureau; some professors in related fields were also included. The average working life of these experts was 10 years, and they had very good professionalism in the field of health emergency. The value of authority coefficient is 0.885 (>0.70), it indicates that the study has a good expert score (22,23,24). In the first round of expert consultation, the coordination coefficient of each item was 0.291(P<0.001), and in the second round of expert consultation, the coordination coefficient was 0.324(P<0.001), which was better than the first round and which indicates that opinions of the experts have agreed. Then, we calculate the mean and variation coefficient (CV) of each concept, and those concepts with a CV>25% and mean<2.5 were deleted (25). The other concepts were modified according to the expert opinion. Finally, combined with the personal interview results, we created a communication framework for influenza pandemic, as shown in Figure 3, that includes the main misunderstandings and the analogies used by the interviewees. The analogies were not coded in our interview. Because this part is to understand what personal experiences or similar concepts respondents will use when understanding the concept of influenza, which may not be familiar to them, so as to provide a reference for risk communication and health education materials. Expert consultation does not involve open questions because they are always able to provide professional advice from a scientific point of view. Figure 3 refers to the format of the FFW model drawn in the study of Lazrus et al. In accordance with the risk factors in Figure 3, the corresponding key concepts are written and were sent to experts for consulting; thus, the final influenza pandemic communication framework is obtained.

**Fig. 3.**
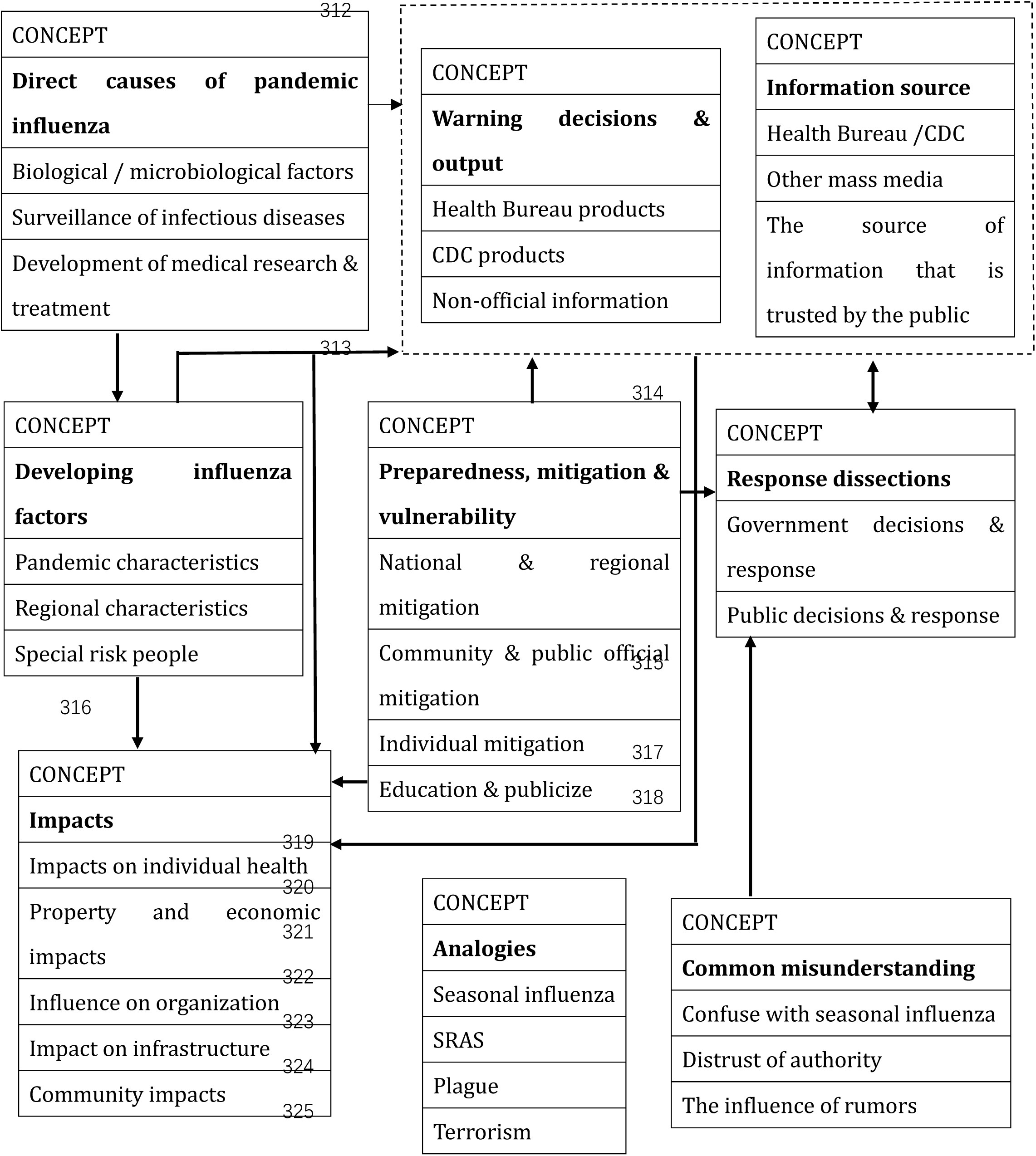
Communication framework of pandemic influenza. The framework is composed of six main conceptual dimensions, the main concept is the bold label and the 2nd-level concept in the box is the part. More detailed concepts in the framework are omitted, see the coding manual in appendix. The whole framework contains 79 concepts, and the arrowhead represents the influence relationship of each part. The analogy part is listed separately to describe the events associated with the interviewees.

### 2. Ruslts of mental model interview

#### 2.1 Overall situation

The percentage of interviewees that mentioned a concept item was counted by the researchers. In addition, this study used a stacked bar chart to show the number of concepts mentioned by the 28 interviewees (Figure 3). As shown in the graph, we distinguish the concept of different attributes in terms of dimensions (risk factors). The depth of the mental model of each interviewee (the number of concepts mentioned by an interviewee) can be visually distinguished by the richness of the color, and we can see which dimensions of the expert’s risk perception the public is highly aware of and in which areas the public lacks awareness. Furthermore, the length of the bar graph reflects the number of concepts mentioned in the dimension: a taller bar graph reflects more relevant concept items mentioned by the interviewees and a deeper degree of understanding of the related content. For example, interviewees 12, 16 and 21 knew more about the emergency response decisions during the pandemic, whereas interviewee #24 was less aware in this regard.

Figure 4 reveals the differences in thinking about the risk of and coping with the influenza pandemic among different groups. Even with a higher education level, each college student interviewee displayed a significant difference in the depth and detail of their mental model. Some of the interviewees’ mental models appear particularly “scarce” (such as interviewee #2 and 25). Almost all interviewees discussed less information than the risk perception of experts. Only one interviewee (interviewee #9) mentioned concepts that reflected almost all parts of the communication framework in Figure 2. The other students did not put forward much more new concepts in the interview. Their conceptual descriptions reflect the concern for specific content and common cognitive deficiencies and misunderstandings. The following sections discuss these most outstanding features of the interview answers.

**Fig. 4.**
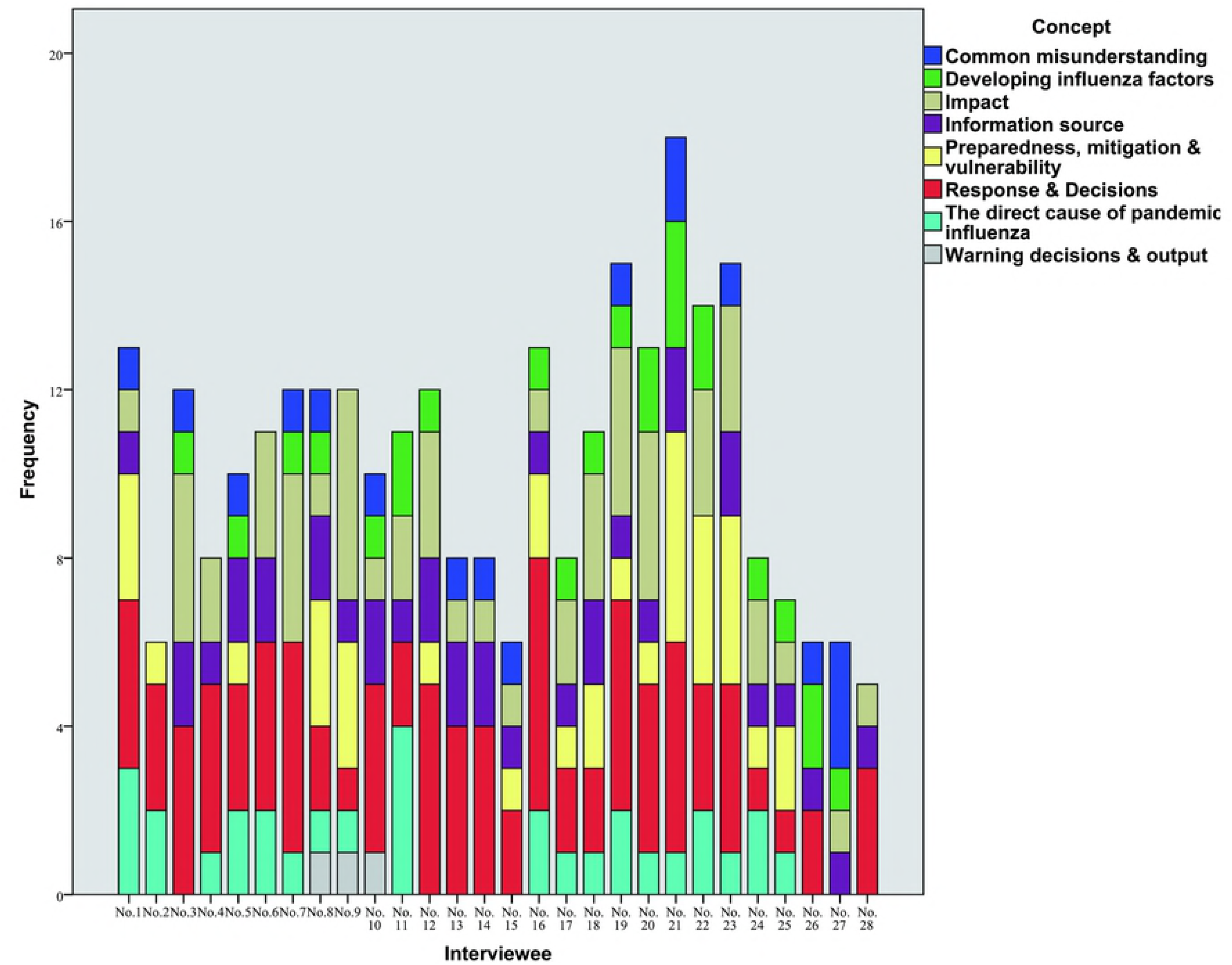
variability in number of concepts mentioned by different interviewee. The straight bars in the graph represent the number of concepts in Fig.3 that can be described by each interviewee. Different colors represent the corresponding conceptual categories. The more colors a respondent has in the bar, the higher the range of understanding of Fig.2 framework he may has.

### 2.2 The formation and development of the pandemic

The interaction of multiple factors may affect the formation and development of pandemic influenza. Some factors are shown in Table 2, 39% of the interviewees believed that influenza virus variation was an important cause of the pandemic. These interviewees used statements such as “new virus”, “virus mutation”, “an unknown virus”, etc. Additionally, 32% of the interviewees referred to disease surveillance, which included “*poor supervision of the source of infection*” and “unchecked work”, and they were more inclined to use terms to express their views (for example, “gene mutation”, “isolation treatment”, “infrared surveillance”, and “take the body temperature”). 46% of the interviewees mentioned characteristics related to the international spread of the pandemic. Interviewee #6 mentioned: “*foreign virus carriers from foreign places into Beijing*”, and interviewee # 9 mentioned “*many are due to alien invasive species, or foreign people bring them to our country*.” However, there are some interviewees believed that climate factors could lead to flu cases because they confused pandemic influenza with seasonal flu, such as interviewee #7, who answered “*when the seasons change, people catch cold easily and catch cold. If they do not pay attention, a pandemic will happen if they don’t do that.*” Many interviewees (46%) also mentioned the impact of population density, including densely populated places and more floating cities with higher risk areas for influenza. For example, interviewee #11 said, “*first of all, (there) is an infected person, and if it is not checked properly, then the person may go to some crowded shopping malls, public places, or schools, and there may be a lot of similar cases in the next few days.*” Interviewee #17 stated that “*… my hometown, a small city, but if it happens, it may take strict measures like the SARS epidemic.*” Other factors that were mentioned less frequently (by less than 17% of the interviewees) included virus resistance, viral power, avian influenza immunity, and a human lack of immunity to new viruses.

**Table 2.** The discussion of related items.

| Concept | Frequency | Ratio |
| --- | --- | --- |
| Virus mutation | 11 | 39% |
| Virus infectious force | 5 | 17% |
| Drug resistance of virus | 4 | 14% |
| Vaccination & related research | 2 | 7% |
| Disease surveillance network | 9 | 32% |
| Lack of immunity | 4 | 14% |
| International dissemination | 9 | 32% |
| Mobile population density | 13 | 46% |
| Health emergency capacity | 4 | 14% |

In addition, the interviewees also discussed the process and development of influenza in light of many factors, some of which were described in greater detail. For example, interviewee 12 described the following situation: “*In one or two days, 3-4 patients with the same symptoms came to the hospital, and then more and more patients will emerge… News will also come out to report, what public places have been taken by some carriers, and what have been done, such as those that came before planes. The events may become clearer and clearer. After that, there may be government departments coming out, some routine inspections that are usually not available, and your units or schools will take some actions.*” However, some of the interviewees also gave a fairly brief overview, stating that “*pandemic influenza is the result of a new virus, the result of the continuous flow and spread of the infected people without control*” (interviewee 3).

Compared to the experts, the mental models of many of the students interviewed contained only part of the communication framework. Although some key elements were mentioned by most of the interviewees, other important factors were rarely mentioned or misunderstood by the interviewees. For example, interviewee #16 believed that the flu was “*foodborne diseases*” and “*caused by drugs*”. For individuals infected with influenza, no one discussed the impact of some vulnerable groups on the development of the pandemic, and there was no further detailed description of the virus variation. Although people do not need to know the details of all the risk factors related to pandemic influenzas, these items present in the communication framework are still an important cause of the pandemic influenza. A full understanding of this information can help people to evaluate risk level in the environment, including which situations may lead to infectious diseases and where there is a greater risk of epidemics.

### 2.3 The impact and consequences of a pandemic

As for the cognitive aspect of the pandemic, the interviewees mentioned the many possible effects of influenza. However, in these general categories, there were some differences in the impact of influenza mentioned among the interviewees. As shown in Table 3, approximately 29% of the interviewees discussed the fatality of the flu, but only 14% of the people described the serious symptoms that could occur after the infection, such as interviewee #5: “…*if there goes a pandemic, it would be more than common cold. Runny nose and sneezing or, maybe, pneumonia?*” None of the interviewees mentioned any complications related to influenza infection. Even if a real pandemic is only comprised of common symptoms of fever and fatigue, complications such as pneumonia, myocarditis, bronchitis, etc. are the real causes of death in some vulnerable patients (26) (27). Therefore, although most of the interviewees understood that the flu can have serious health threats, they did not understand how people die as a result of the flu. These misunderstandings may be related to some interviewees’ subjective and one-sided understanding of the pandemic and the lack of targeted health education. For example, interviewee #10 said, “*That is, people usually don’t pay attention to clothes, then they catch a cold. It is quite a normal situation every year*.”

**Table 3.**
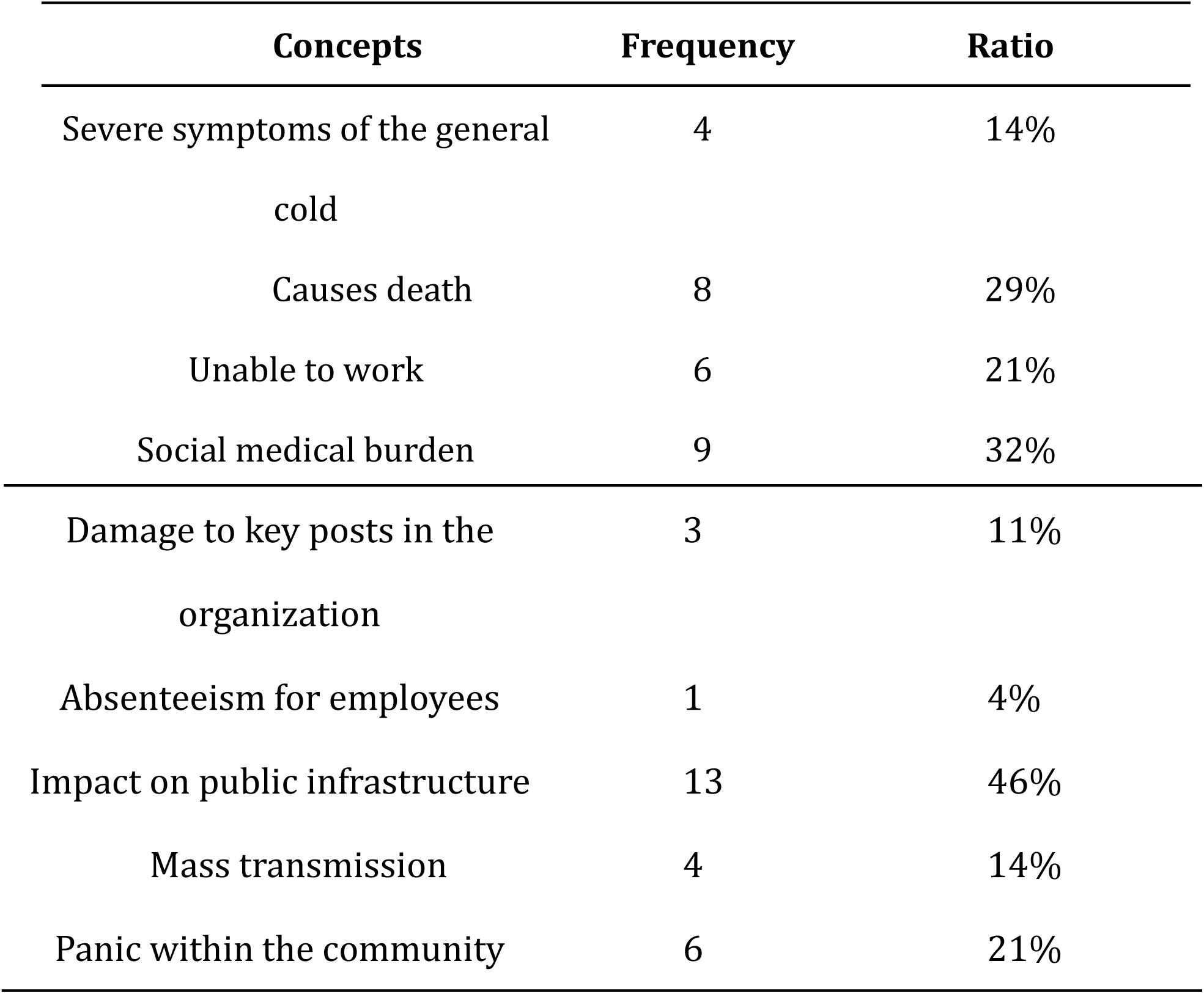
The discussion of related items.

| Concepts | Frequency | Ratio |
| --- | --- | --- |
| Severe symptoms of the general cold | 4 | 14% |
| Causes death | 8 | 29% |
| Unable to work | 6 | 21% |
| Social medical burden | 9 | 32% |
| Damage to key posts in the<br>organization | 3 | 11% |
| Absenteeism for employees | 1 | 4% |
| Impact on public infrastructure | 13 | 46% |
| Mass transmission | 4 | 14% |
| Panic within the community | 6 | 21% |

Most interviewees also discussed the social and economic impact of the pandemic, and 46% of the interviewees referred to the negative impact of schools, shops, public transport and other infrastructure during the pandemic, such as interviewee #14: “*schools may shut down… The shops outside may be closed because of this disease, and the economy may be seriously affected because everyone will hide at home.*” Most of the types of infrastructure, of which transportation was mentioned the most frequently, were generally quoted as examples of people during the SARS or bird flu period, such as interviewee #11 stated: “*everyone is not going out at the time of the outbreak…wearing a mask if you have to go.*” 32% of the interviewees were worried about overburdened hospital patients during the pandemic. Some of the interviewees (28%) have also been associated with disastrous consequences, including the impact of the pandemic on the community, suggesting that they recognize the possibility of a broader and indirect impact of the pandemic. According to interviewee #16, “*For a long time… Our life may be threatened, many people steal food, drugs, and be locked inside their house…not just the direct impact, it will definitely bring other serious problems.*” Although the relevant concepts in the communication framework were mentioned, the interviewees pay more attention to consider about the possible impact of a pandemic on the organization or community, they fail to understand the serious damage that pandemic influenza could cause to individual health, nor is it fully aware of the characteristics of the pandemic itself in all its periods.

Surprisingly, 30% of the interviewees believed that a negative impact of a flu pandemic would be minimal or more positive, and almost all of them said, “*fells like the pandemic is far away from me.*” According to interviewee #23, “*Is a kind of epidemic disease, but speaking of cold and flu, what is generally not a major disease, easier to treat the feeling, plus the pandemic, it is only a larger scope of infection, right?*” Moreover, “i*t’s not a problem to suffer a pandemic… the SARS also created a great electric business, maybe, the world needs to be rearranged*” (interviewee #22). The content reflects that some students still do not pay much attention to public health and their own health. More people choose to passively wait and accept the strategies and measures taken by the school or the state government; they lack the initiative to understand the relevant information and take preventive actions. In addition, the lack of targeted publicity and education may also be a reason for the blind optimism of these interviewees. For example, interviewee 3 stated that “*(it will) find a way to deal with it at last. I don’t care much about this. I’m young and strong after all.”*

### 2.4 The countermeasures of the pandemic

Coping strategies in Table 4 are essential to pandemic emergency work and an important part of the communication framework in Figure 2. 29% of interviewees mentioned the importance of personal hygiene habits, such as wearing masks and isolating patients, but there are not many people who provided detail regarding these aspects. A few interviewees described these strategies on the government, organization, and individual levels. Most of them referred to “*masks*” and “*be far away from the cough*” in the relevant open description, and mentioned details of whether to use a special mask or separate the patient from the family, etc. There was no mention of personal protective measures. For example, interviewee #6 said: “*if it is more serious, wear a mask, and then the hospital will be more nervous about the flu, and there should be nothing else.*” Another 18% of the interviewees believed that there was no need to isolate the suspected patients, such as interviewee #19: “*You cannot go to the hospital first, because most of the cases are not true flu, to the hospital may really be isolated, so look first.*” For the government’s decision-making, 57% of the interviewees mentioned health education and counseling. Most of them willing to accepted the necessary emergency response; over 1/4 of the interviewees referred to influenza surveillance, public disinfection, and hospital treatment. These answers demonstrate that the students with better educational backgrounds have a certain understanding of the government’s coping strategies and have a high degree of potential coordination, but there were some mistakes and lack of understanding in the concept of the most effective protection decisions that an individual can make. However, although vaccination is the most effective way to prevent flu, only 2 of the interviewees said they were willing to receive the flu vaccine, and the other interviewees said they would not vaccinate themselves if they were not compulsory. “*There is no need for voluntary vaccination*” (interviewees 3 and 17), “*some vaccines may have side effects…it will hurt me.*” (interviewee 26) and “*the vaccine seems to prevent ordinary flu, and there is no special vaccine for bird flu.*” (interviewee #12).

**Table 4.**
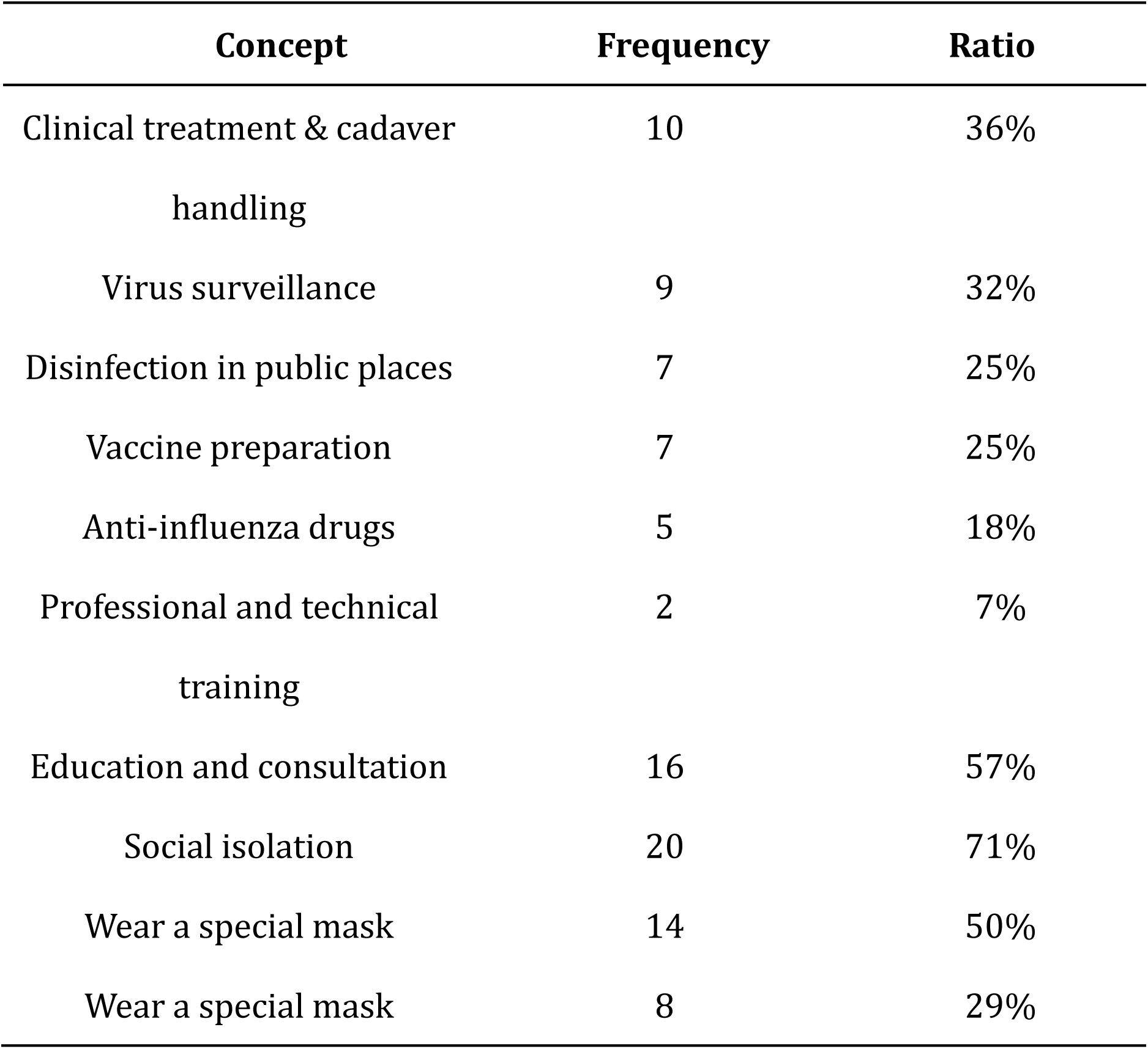
The discussion of related items.

| Concept | Frequency | Ratio |
| --- | --- | --- |
| Clinical treatment & cadaver handling | 10 | 36% |
| Virus surveillance | 9 | 32% |
| Disinfection in public places | 7 | 25% |
| Vaccine preparation | 7 | 25% |
| Anti-influenza drugs | 5 | 18% |
| Professional and technical training | 2 | 7% |
| Education and consultation | 16 | 57% |
| Social isolation | 20 | 71% |
| Wear a special mask | 14 | 50% |
| Wear a special mask | 8 | 29% |

Notably, interviewee #9. who come from Hong Kong, were able to relatively fully describe the individual and government contingency strategies and discussed their own personal experience of avian influenza in Hong Kong in addition to the elaborating on the entire process of emergency work. For example, he mentioned, “*… all the rumors about poultry industry will quickly corrected, and poultry killing is very efficient*. *The response of the government’s Department (Ministry of prevention) is very sound, and we know how to check the body temperature, wash hands, and a lot of protection will be done. We have confidence in the epidemic prevention system of Hong Kong.*” He also stressed that “*the individual should understand the knowledge of influenza*”, and “*to understand the extent of the risk of their own ar*ea.”

This response fully embodies the maturity and perfection of the Hong Kong government in the risk communication of emergencies and the higher risk awareness and compliance of the compatriots in Hong Kong; moreover, this interviewee has sufficient understanding of the government’s emergency disposal process, the measures that can be taken and how the people can help; thus, the related communication and publicity strategies are worth to refer.

### 2.5 The acquisition of risk information and public suggestion

The risk of pandemic influenza can be reduced by timely warning, access to correct information, and attitude toward communication and interest in the face of threats. As shown in Table 5, 50% of the interviewees had a certain information identification ability; 43% of the interviewees chose to obtain their information on pandemic risk from the official channels. All the interviewees were willing to take several methods to search for risk information including using the Internet. In addition, a considerable proportion of interviewees would also choose other information access methods for personal preference, such as “go to the library to consult the literature” (interviewee 1) and “*ask my doctor friend*” (11). As a whole, the students’ trust in the government was generally high. In case of emergencies, they would first turn to the authority of the government and would choose a variety of ways to obtain relevant information on the pandemic to help themselves to make decisions. However, although first-hand influenza warning and decision support information come from the CDC, very few of the interviewees (10%) were able to clarify what types of communicators can provide help and provide detailed description on this topic, including what types of early warning information are available, where the information is, and how the information is transmitted. People have only mastered the general concept, such as interviewee #11: “…*go to the official website or WeChat (to) find how to prevent*.” As for health education and publicity, most of the interviewees said that they would not take the initiative to participate in similar activities. The reasons were “*traditional lectures are boring*”, “*the publicity manual was not attractive*”, etc. And it just like interviewee #25 said, “*I think all of them are theoretical knowledge, and you are also the same on the Internet. If they can tell us something that you need to deal with when an event comes, it would be better.*”

**Table 5.**
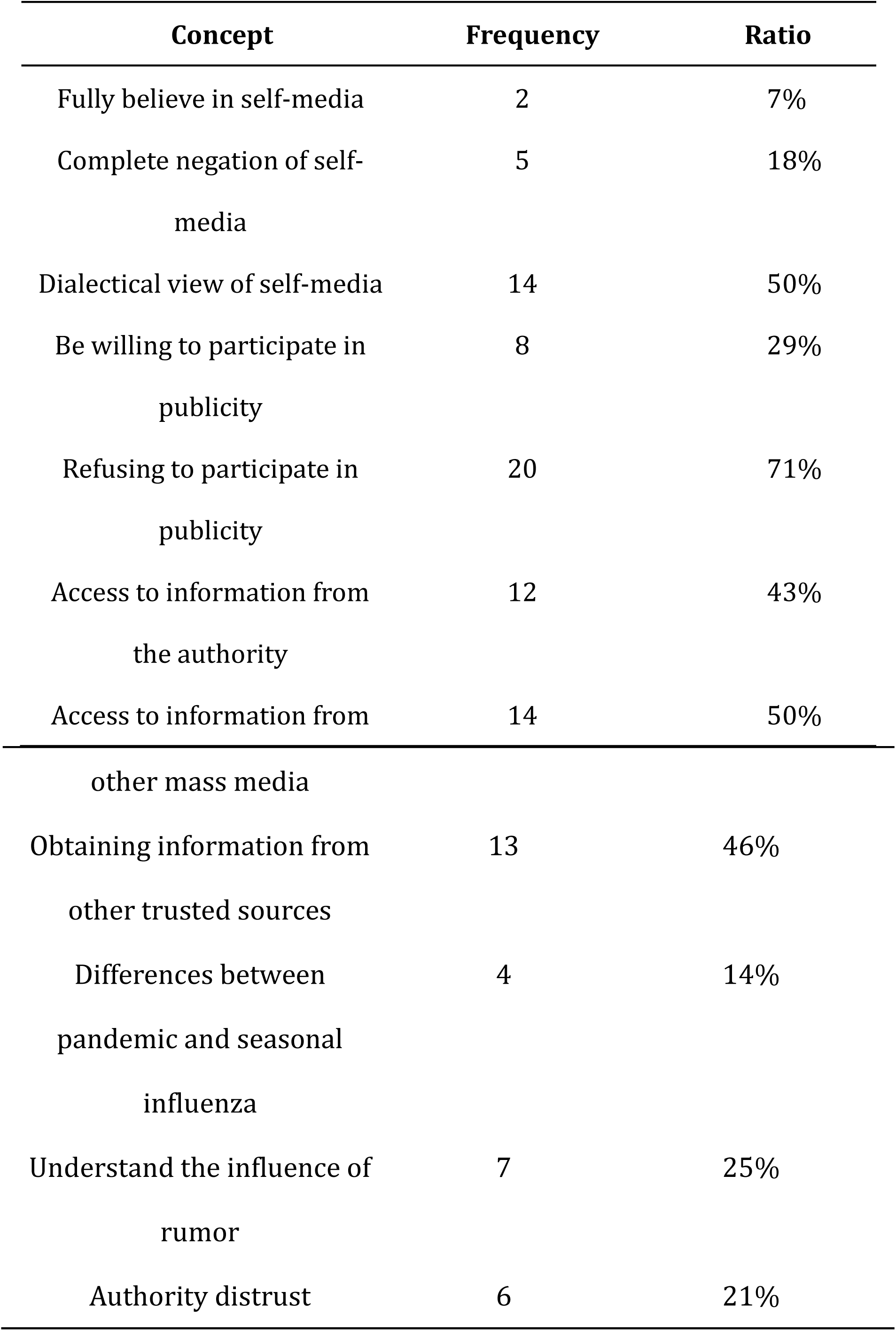
The discussion of related items.

At the end of the interview, the researchers asked the interviewees for suggestions regarding future risk communication. Most of the interviewees were satisfied with the current government’s work and have positive attitude towards the emergency plan of the official guide form, they were more focused on “*the details of the emergency work*” (mentioned by 25% of the interviewees) and “*hope to get official plan*” (mentioned by 21% of the interviewees). For example, interviewee 20 says that: “*…the way must be change, not as before, because the flu is not like a normal cold, people will not pay much attention to it. Communication, whether it is a family or school, it is best to have some specific suggestions, such as how to wash hands and disinfection, everyone can refer to themselves to do it.”*

### 2.6 Interviewees’ personal experience and analogies

During the interview, the interviewees showed certain characteristics on the macro concept of pandemic influenza. A total of 32% of the interviewees cited “SARS” and “avian flu”, for example, interviewee# 9 said, “*the disease like H1N1, it should be a pandemic*.” “*Similar to it, there will be less people in the street, the economy will be paralyzed, the hospital will be slower and people are confused.*” Additionally, 17% of the interviewees compared a flu pandemic to the “Black Death” during the history of Europe, such as interviewee 23: “T*he black death, the plague and so on. At that time, leprosy established leprosy villages. On a large scale, I think SARS is not big enough. The pandemic that I define here will be larger and more harmful.*” Most interviewees were more connected and described the common cold or seasonal flu, with a limited descriptive vocabulary. For example, interviewee #17 stated that “*the pandemic is possible, the flu, and many people may be involved, which is involved, but first of all these people are the common cold.*” 17% of the interviewees mentioned the effects of haze pollution and environmental damage, which were not involved in our communication framework. Another 2 interviewees believed that the pandemic influenza was a “*terrorist attack*” and believed that a pandemic was uncontrollable, for example, “*the flu is unavoidable*” (interviewee #28) or “*someone destined to get sick*” (interviewee #12). Interviewee #9 described pandemic influenza as “the decay of the human’s mind - if we do not treat the people and things around us, our evil will bring flu, and that is the retribution we have suffered”; this view was a result of his religious beliefs.

These statements illustrate how personal experience affects mental models and their environment. In the absence of influenza knowledge and other directly related knowledge, interviewees may try to combine their understanding with all available personal experiences to fill the gaps in the information. As the research of Visscher et al indicates, the actions taken by people in the face of unknown risks are first based on relatively familiar personal experiences, the impact of risk media coverage and common-sense education, which are often used to estimate the severity and consequences of events and to take into account the impact of unknown risks and consequences (28). However, exaggerated associations or analogies can be misleading in interpreting risk factors and coping strategies. For example, in the study of flash floods risk communication by Lazrus et al., some of the interviewees planned to provide flood control sandbags for each family to make every effort to block the flood outside their house, after viewing the photos of Hurricane Katrina (16). The over-speculation of emergencies may result in negative effects on real threats, including the stealing of daily necessities and vinegar, which occurred in the SARS period (mentioned by 7 of the interviewees).

## Discussion

In general, there was a clear difference in the breadth and depth of the overall understanding of the pandemic-related information and communication framework among each student interviewed. As expected, in the context of communication framework, most of the students’ mental models were not as rich as those of the experts, and they were more concerned with the key information necessary to make individual decisions in the interpretation of risk information (i.e., when to act and in the form of the pandemic: “Now, what kind of impact will it cause, and what kind of protection measures can protect me?”) Most public interviewees only referred to the key concepts in the communication framework, but without a detailed description, or in an inaccurate or unclear manner; therefore, these gaps may reduce the ability of people to manage their own behavior and their compliance with expert opinions. Compared to the communication framework in Figure 2, the interviewees used personal experience and analogies to produce more related concepts to establish the information base they needed to make decisions.

### 3.1 The causes of misunderstandings

As soon as the human body get infected by flu virus, the related symptoms includes high fever (up to 39 degrees centigrade or higher), muscle aches, etc., accompanied by dry cough and severe fatigue, and complications in the heart and lung systems, particularly in those with low immunity. Among the elderly, infants and young children, this is also an important cause of potential lethality (29). The interview results show that some students do not pay enough attention to the impact of pandemic influenza and remain blind and optimistic, particularly regarding its potential lethality, serious complications, and the identification of vulnerable populations. The trust in the country’s sound epidemic prevention system, the desalination of the history of the epidemic, and the lack of targeted health education are the main reasons for the over-optimism of the interviewees. Consequently, the students who have the wrong risk perception will estimate themselves as “strong young people” or “enough understanding of the flu”. Once the flu is outbreak, it may also bring misleading information to other individuals in their social circle, which will affect their emergency decisions. In particular, for those who have experienced influenza pandemic without being negatively affected, luck may cause them to have a more positive response to future pandemics (30).

Secondly, although the H1N1, H5N1, and other influenza outbreaks have been derived from the new virus from mutation, the repetition of the old virus and the prevention of the flu season still risk becoming a pandemic (26). Therefore, for college students, paying attention to the prevention of common seasonal influenza and simultaneously being able to distinguish the key differences from the pandemic can effectively improve the level of personal risk cognition. Among the interviewees, we found that some students were still confused about the concepts of pandemic and seasonal influenza: they believed that a pandemic is the mass spread of seasonal influenza or that a pandemic is an almost impossible “super calamity”, when in fact a pandemic cannot be completely avoided. Moreover, a pandemic is often unpredictable and generally involves international diffusion. Therefore, it is important to understand that the pandemic is not far away from us. We must pay attention to our own prevention during the flu season and have a certain understanding of the symptoms of an abnormal cold, particularly when traveling abroad; otherwise, the patient may mistake their symptoms for a common flu. Medication has delayed the timing of diagnosis and treatment, resulting in serious consequences.

Finally, about the vaccination, our interviewees have negative views about that. Only 2 of the 28 interviewees mentioned the importance of the vaccine and had a history of active vaccination, and the reasons mainly focused on the conventional “I feel good and don’t need vaccination”, “not knowing about the importance of vaccine”, and “Doubts about the safety of vaccines”. In fact, the extremely low vaccination rate of influenza vaccines in China, the low vaccination rate is a long-standing problem. Therefore, our risk communication at present seems inadequate in promoting the necessity of vaccination, and the public is not aware of the importance of the vaccine for influenza prevention or the misperceptions caused by its one-sided understanding of the pandemic, as discussed in 2.4. In the investigation of the willingness of the elderly to be vaccinated, Geng Shaoliang (31) found that the main sources of influenza and related knowledge in the elderly were family, relatives, friends and television, and the most trusted means of knowledge were doctors. There are cracks in clinical and public health knowledge, and clinicians lack knowledge on the importance of vaccination. Many doctors have reservations regarding vaccinating the elderly, and the rate of influenza vaccination for medical staff is not sufficiently high. Therefore, the correction of this misunderstanding is important for both college students and because it can promote the dissemination of inoculation knowledge of young students in the family, thus promoting the inoculation rate of the recommended group (the elderly and young children).

### 4.2 The defect of individual mental model

As discussed in section 2.5, in the absence of relevant knowledge and information, the interviewees applied personal experiences and analogies to make up the foundation of their mental model and to help themselves understand the risk of the pandemic. Understanding differences in causality between risk factors can also lead to important differences in risk perception and coping between individuals (32). Many students only know a few general concepts and have not yet formed a complete emergency preparedness mode of thinking in communication framework, knowing what you can do during the pandemic, but not much about what to do and what is truly meaningful. For example, although almost all interviewees mentioned wearing masks and bringing in patients in time for medical treatment, the most basic measures can be limited in the presence of a real pandemic, which is only a result of a personal experience analogy (compared to a cold or related disease). The expert opinion indicates that in the period of the pandemic, suspected patients should first undergo home isolation observation to avoid causing infection while receiving medical treatment or engaging in activities, and relatives and friends should avoid contact with them as much as possible. In addition, most interviewees have only basic concepts (the government, the health department) regarding the types of communicators, who provides the relevant risk information, how to access these channels, etc. These overly broad understandings may limit their ability to identify critical information quickly or influence their knowledge of certain information under the threat of serious flu, particularly when their usual sources of information or communication channels are not available or the necessary information is not provided, such as the presence of a new influenza virus. In a short period of time, officials are unable to give exact messages or be out of protection from the spread of information. Public trust in official authority may be reduced.

### 4.3 The traditional publicity requires improvement

As a young generation, college students enjoy new things, with the rapid and diversified development of the current social information dissemination mode. Therefore, emergency workers must change the form and content of influenza health education. Students prefer health education with new forms and systematic content. The appeal of traditional lectures and guideline book full of scientific word is far less attractive, and it is hoped that the government will “reduce the over generality of the description” and “release relevant data to increase persuasion” in future communication work. Foltz’s research confirms that it is necessary to use a variety of mechanisms in the risk communication of emergencies. People with nonprofessional backgrounds tend to think in more specific terms, their vocabulary is less expansive, and subtle expressions cannot be well understood. They are easily attracted by bright colors and charts. Complex text information transmission will make people feel tired and irritable (33). If possible, 2 student interviewees also suggested that students can organize practical exercises, such as self-sterilizing in family homes or dorms, which they think is more helpful to deepen the impression and understand some of the self-protection measures used to cope with the pandemic.

For advice on pandemic influenza health education and risk communication, we should let students know which government departments can help them. The study results of Garrett et al. show that information consistency is the decisive factor in understanding and perceive personal risk. In terms of communication effectiveness, multiple sources of consistent messages are usually more effective than messages from a single source or with different contents. The earlier the warning people receive and the greater the threat of information, the greater the possibility that people take active preventive measures (34). Based on the results of this study, the students have a certain ability to identify unofficial risk information. The probability of being affected by rumors at the early stage of the pandemic was not significant. If the official information publishers of the pandemic are able to understand them, they can obtain relevant information according to their own needs and improve their official information, creating a degree of trust in the authority. Therefore, the government department can refer to the communication content proposed by the research institute to incorporate the risk information required by the student group into influenza warnings and information communication, including the relevant situational information and the proposed measures, and maintain the consistency of multiple communication messages. These messages can be used not only for the notification of influenza warnings but also for the warning of influenza to provide support for follow-up long-term public health education.

## Conclusion and Proposal

As an emergency communicator, the goal of government is to integrate the expertise of medicine, epidemiology, behavior, and statistics into the information and concepts that can be understood by the public. Although the current related studies are mostly descriptive, the quantitative evidence-based assessment of the risk communication of emergencies may occur with the standardization of the practice and the expansion of the ability to determine whether the message is transmitted to the designated group and the extent to which the population will apply and understand the information. After SARS, China put significant attention on the prevention and control of large-scale infectious diseases. However, studies of risk communication remind improved. Through personal interviews, our researchers explored the “risk cognition boundary” of Beijing college students regarding pandemic influenza and clarified how students understand information on the pandemic and what they are most concerned about. Although they may know a bit about the SARS epidemic in history, there are many problems regarding the potential risk of pandemic influenza in the future, which our communicators must pay attention to and solve.

Unlike many other catastrophic events, the pandemic influenza does not directly affect the normal operation of university. Although the related hardware facilities will not be damaged by the disaster. College students, who leave home for a long time to live independently, will be hard to continue with their studies and take care of their lives simultaneously under the threat of sudden flu outbreak. Therefore, a college pandemic influenza plan is necessary. In addition, as a risk communicator, the government department releases related risk information on pandemic influenza in the form of a reference plan. It can effectively bridge the gap between the public and experts on the understanding of influenza risk information and ensure the effectiveness of communication for the long and short term.

According to the study result we suggested that pandemic risk communication should focus on the following aspects: ① influenza virus variation (particularly the influenza A virus) and seasonal influenza have the potential to evolve into a pandemic, and the prevention of common influenza cannot be ignored; ②the impact of a pandemic is often unprecedented; influenza virus infection can be fatal, but it also results in serious complications in addition to severe cold symptoms; ③ influenza vaccination has a positive effect on the prevention of a pandemic and should be actively administered, particularly in young children with low immunity and elderly people; and ④ for suspected patients, the first preventative measure is home isolation, and the protection of the family members is important because it is very dangerous to keep in close contact with them. The key to improving the ability to handle public health emergencies is to master the common sense of influenza preparation and the correct personal response and to understand the degree of risk in the region and the individual’s pandemic response plan. In addition, to help people understand information and enhance the compliance of certain coping strategies, we can try to provide some “rules of thumb” (35) from experts to help the public to infer the conclusion of the unknown disease. Moreover, the local health and epidemic prevention department, as an authoritative communicator and decision-maker, can also give schools certain support for health care during the pandemic, such as the use of the network for remote health education or local radio or television stations to guide the students to respond to the work.

In order to solve the key gap in public understanding of the risk of pandemic influenza and the response to protection. In emergencies, people can observe the reality of the epidemic but do not know how to find more information about warnings. It is necessary to apply various ways and mechanisms to run publicity. We suggest that the government strengthen the application of new media such as micro-blog and WeChat to adapt to the preference of information acquisition for young people and release early warning information quickly and promptly. Second, in the form of publicity, traditional lectures can be gradually changed to new formats, such as public service films, songs, and situational construction experience. These examples can apply visual and auditory stimuli, not only to provide a new method of communication to the audience but also to avoid the misunderstanding of the information. Scenes created by a video can enhance the personal experience and facilitate the referencing of the analogy to the individual in a risk event, allowing them to make the right risk assessment and response behavior.

## Acknowledgements

The authors would like to acknowledge Linxian Wang for helping compiling interview questionnaires, making suggestions on interview skills and finding supporting documents and express the sincere gratitude to students involved in the interviews of this research.

